# Wheel running increases hyperthermia and mortality rate following 3,4-methylenedioxymethamphetamine (MDMA) in rats

**DOI:** 10.1101/126706

**Authors:** M. A. Taffe

**Author notes:** Address Correspondence to: Dr. Michael A. Taffe, Department of Neuroscience, SP30-2400; 10550 North Torrey Pines Road; The Scripps Research Institute, La Jolla, CA 92037; USA.

## Abstract

Hyperthermic responses are commonly reported in cases of human medical emergency following recreational use of 3,4-methylenedioxymethamphetamine (MDMA, “Ecstasy”), but a precise determination of contributing environmental factors has been elusive given the relative scarcity of threatening and fatal reactions in humans. This study was conducted to determine if elevated physical activity contributes to hyperthermic responses to MDMA in a well controlled animal model. Unrestrained male Wistar rats were monitored with minimally-invasive radiotelemetry techniques following challenge with MDMA (1.0, 5.6 and 10.0 mg/kg, s.c.). Studies were conducted in low (23-25°C) and high (27°C) ambient temperature (T_A_), with and without access to an activity wheel. The study confirmed dose dependent effects on body temperature, chamber activity and wheel activity which were modified by different T_A_ conditions. Increases in wheel and home chamber activity produced by 10 mg/kg MDMA increased the magnitude of hyperthermia under 27°C T_A_. Furthermore, greater subject mortality was observed in the wheel-access condition compared with the no-wheel condition. These data provide direct evidence that sustained physical activity increases the hyperthermic response to MDMA and that this is associated with increased lethality. This is the first direct experimental confirmation that increased physical activity may be a risk factor for adverse reactions to MDMA in human user populations.

## 1. INTRODUCTION

It is well established that disruption of thermoregulation following the administration of 3,4-methylenedioxymethamphetamine (MDMA, “Ecstasy”) occurs in a variety of laboratory species including monkeys (Banks et al. 2007; Taffe 2012 ; Taffe et al. 2006; Von Huben et al. 2007), pigs (Fiege et al. 2003; Rosa-Neto et al. 2004), rabbits (Pedersen and Blessing 2001), guinea pigs (Saadat et al. 2004), rats (Brown and Kiyatkin 2004; Dafters 1994; Malberg and Seiden 1998) and mice (Carvalho et al. 2002; Fantegrossi et al. 2003). As such, it is possible to use animal models to explore the causes of the relatively infrequent, but severe, MDMA-related human medical emergency and/or death (Mascola et al. 2010)in which hyperthermia is a common observation (Dams et al. 2003; Gillman 1997; Greene et al. 2003; Mallick and Bodenham 1997). It is frequently hypothesized (Parrott 2004) that prolonged physical activity (e.g., dancing) in the rave or club environment interacts with the elevated ambient temperature (T_A_) to increase risk of medical emergency but this possibility has not been directly determined in controlled laboratory models. Given that MDMA use patterns are more episodic than patterns for traditional stimulants, and that users are motivated to use scientific results to reduce harm, provision of actionable real-world risk information is likely to result in behavioral change (Taffe 2015). This makes it imperative to rule physical activity in or out as a contributing risk factor and to determine if it interacts with elevated T_A_.

MDMA increases open field locomotion in rats at doses of about 10 mg/kg of either the racemic mixture or the S(+) stereoisomer (Gold and Koob 1988; Gold et al. 1988). MDMA may also increase rats’ locomotion under certain conditions following doses of 1-3 mg/kg (Baumann et al. 2012; Kehr et al. 2011). Unlike open field activity, the access to an activity wheel can be varied without undue restraint and therefore it can be used as an independent variable to model the presence or absence of sustained physical activity (i.e., the dancing that is a feature of one prominent Ecstasy use environment). In rats, the thermoregulatory effects of MDMA depend on ambient temperature (T_A_) with a *decrease* in body temperature observed at a sufficiently low (∼20-23°C) T_A_ and an *increase* in body temperature at elevated (∼27-30°C) T_A_ (Dafters 1994; Malberg and Seiden 1998). The MDMA dose tends to overlay, rather than adding to or potentating, the effect of T_A_ since higher doses can produce *hyper*thermia under lower T_A_ conditions where lower MDMA doses result in hypothermia. Low T_A_ is also protective against MDMA-induced lethality in both rats and mice (Fantegrossi et al. 2003; Malberg and Seiden 1998). Interestingly MDMA appears to elevate body temperature across a similar T_A_ range in monkeys or humans (Crean et al. 2007; Crean et al. 2006; Freedman et al. 2005; Taffe 2012; Taffe et al. 2006; Von Huben et al. 2007) which suggests that rodent studies conducted at T_A_ which reliably produces *hyper*thermia, i.e., within the thermoneutral range (Gordon 1990), are most relevant to the human condition.

We have shown in prior work that MDMA increases rats’ wheel activity after a 10 mg/kg dose, but decreases wheel activity after 7.5 mg/kg and lower doses, when administered in the dark part of the circadian cycle under normal laboratory T_A_ (∼22-23°C) conditions (Gilpin et al. 2011; Huang et al. 2012). Wheel activity is similarly increased by the prototypical stimulant methamphetamine and the novel cathinone derivative 3,4-methylenedioxypyrovalerone (Huang et al. 2012), which supports the validity of this assay. The Gilpin et al. (2012) study found no difference in the rectal hyperthermia generated by 10 mg/kg MDMA when animals had access to the wheel compared with that observed without wheel access, however an apparent increase in mortality rate was found. Since the thermoregulatory response to MDMA in rats depends on the T_A_ in addition to the dose (Dafters 1994; Malberg and Seiden 1998) the present study was conducted to determine if wheel access increases MDMA-induced hyperthermia under differential ambient temperature conditions. A minimally invasive radiotelemetry system, shown to detect alterations of both the activity and body temperature of rats after administration of methamphetamine, 4-methylmethcathinone, Δ^9^-tetrahydrocannabinol, α-pyrrolidinopentiophenone and 3,4-methylenedioxymethamphetamine (Aarde et al. 2013a; Aarde et al. 2015; Taffe et al. 2015; Wright et al. 2012) (and most specifically 5.6 mg/kg MDMA, s.c. under a T_A_ of 30 °C; (Miller et al. 2013)), was used to reduce potential effects of the stress of the rectal sampling used by Gilpin et al. (2012).

## 2. Materials and Methods

### 2.1 Animals

Male Wistar rats (Charles River, New York) were housed in a humidity and temperature-controlled (22°C ±1) vivarium on a 12 hr : 12 hr reverse light cycle. Animals were 250-300g on arrival at the laboratory. Drug challenge studies were conducted in the animals’ dark (active) period. Animals had *ad libitum* access to food and water throughout the course of the studies. All procedures were conducted under protocols approved by the Institutional Care and Use Committee of The Scripps Research Institute and consistent with the US NIH guidelines for the care and use of laboratory animals (Garber et al. 2011).

### 2.2 Equipment

Activity wheels (Med Associates Model #ENV-046; ∼35 cm diameter runway) were attached to the side of a shoebox style home cage modified with a door to provide wheel access. Heating of the experimental rooms was by individual space heaters and verified by a portable thermometer and a probe attached to the telemetry system (Data Sciences International; C10T). Variability of the ambient temperature was ±1°C from the target temperature for these studies.

### 2.3 Radiotelemetry

Radiotelemetry transmitters (Data Sciences International; CTA-F40, TA-F40) were implanted using sterile surgical techniques under anesthesia with an isoflurane/oxygen vapor mixture (isoflurane 5% induction, 1-3% maintenance). An incision was made along the abdominal midline posterior to the xyphoid space, just large enough to allow passage of the miniature transmitter which was placed in the abdominal cavity. Absorbable sutures were used to close the abdominal muscle incision and the skin incision was closed with the aforementioned tissue adhesive (3M^™^ Vetbond^™^ Tissue Adhesive; 3M St Paul, MN). A minimum of 7 days was allowed for surgical recovery prior to starting the study. For the first three days of the recovery period, cephazolin (0.4 g/ml; 2.0 ml/kg sc; once daily) and flunixin (2.5 mg/ml; 2.0 ml/kg sc; once daily) were administered.

Radiotelemetry recordings of body temperature (°C) and activity (total counts which are derived from the rate recorded by the software multiplied by the number of minutes in the sampling interval) were made every five minutes via a telemetry receiver plate placed under a normal shoebox style lexan cage. In all studies a response plan for managing excessive hyperthermia was in place. If an animal’s temperature exceeded the allowable threshold (42°C) it was placed on a pad covering a layer of ice in a standard chamber until normative temperature range (36.5-39°C) was restored for at least an hour. Animals were then monitored periodically up to ∼6-8 hours after the dosing time. Animals that could not maintain normative body temperature or were observed to be unresponsive, immobile and/or moribund this long after dosing were euthanized as a matter of protocol.

### 2.4 Drugs

The (±)-3,4-methylendioxymethamphetamine HCl for this study was provided by the US National Institute on Drug Abuse Drug Supply program. Drug doses (expressed as the salt) were diluted in physiological saline and injected subcutaneously in a volume of ∼1 ml/kg. Treatment conditions (drug dose, wheel access) were tested in a randomized order across groups with a minimum 3-4 day interval between drug challenges.

### 2.5 Data analysis

Analysis of the temperature, activity and wheel running data employed analysis of variance (ANOVA) with within-subjects factors of time post-injection. The drug treatment condition with and without wheel access factor was analyzed as between-subjects due to differential subject dropout across treatment conditions. The data were averaged (temperature, wheel quarter rotations) or summated (activity counts) across 30 min intervals for analysis. Any significant ANOVA main effects were followed with post-hoc analysis using Holm-Sidak correction for multiple comparisons. The first post-hoc strategy compared the highest dose of MDMA with Wheel access to all other dose/access conditions within experiment to test the main hypothesis under investigation. The secondary post-hoc strategy tested all dose/wheel access conditions against all other conditions. Analyses were conducted using Prism 6 for Windows (v. 6.02; GraphPad Software, Inc, San Diego CA). Graphs were generated with Excel (Microsoft, Redmond WA) and figures created in Canvas (v.12; ACD Systems of America, Inc, Seattle, WA)

### 2.6 Experiments

#### 2.6.1 Experiment 1: Effects of 1, 5.6 mg/kg MDMA at∼23°C Ambient Temperature With and Without Wheel Access

A group of male Wistar rats (N=6; 62 wks of age, mean 630 g) were used in this initial experiment. This group had been used in a prior study of the effects of methamphetamine (1, 5.6 mg/kg s.c., under 20 or 25 °C ambient temperature conditions) as previously reported (Aarde et al. 2013b). Ambient temperature (T_A_) was not manipulated recorded but was estimated as about 23 °C based on the building / facilities thermostat settings. The main goal was to validate the model telemetric monitoring of MDMA responses in the presence and absence of wheel access, as a technical improvement over our prior study (Gilpin et al. 2011), under conditions unlikely to produce threatening hyperthermia. Doses of 0.0, 1.0 and 5.6 mg/kg, s.c., were evaluated with and without wheel access in a randomized order.

#### 2.6.2 Experiment 2: Effects of 1,5.6 mg/kg MDMA at 25°C Ambient Temperature, and 10 mg/kg MDMA at 23°C Ambient Temperature, With and Without Wheel Access

A group of male Wistar (N=8; 20 wks of age, mean 488 g body weight at start) were used in these studies. Animals had no drug exposure prior to the start of this study. After first completing the mixed order challenges of 0.0, 1.0, 5.6 mg/kg MDMA s.c. with wheel and no-wheel access conditions under 25 °C, the group was next used to evaluate 0.0 vs 10 mg/kg MDMA s.c. with wheel/no-wheel access conditions in another randomized order, repeated measures design under 23 °C.

#### 2.6.3 Experiment 3: Effects of 5.6, 10 mg/kg MDMA at 27°C Ambient Temperature, With and Without Wheel Access

Two different cohorts of male Wistar rats were used, including N=10 ∼40 wk old animals and N=6 ∼20 wk old animals. The latter group was drug naïve. Subsets of the former group previously participated in the Experiment 2 studies and in pilot investigations of the effects of 5.6 mg/kg MDMA and 5.6 mg/kg MA at 20 °C T_A_ as well as 5.6 mg/kg (+)-MDMA vs 5.6 mg/kg (−)-MDMA at 27 °C T_A_ (N=2). Another subset received prior exposure to 1.0, 5.6 mg/kg 3,4-methylenedioxyamphetamine 23 °C T_A_ (N=2). For this study the individuals received 0.0, 5.6 or 10.0 mg/kg MDMA s.c. with wheel and no-wheel access conditions in a randomized order under 27 °C ambient.

## 3. Results

### 3.1 Mortality

The 16 rats treated with 10 mg/kg under 27 °C T_A_ in the wheel condition (Experiment 3) consisted of 10 from the ∼40 week old group and 6 from the ∼20 week old group; of these, 6 of the older rats and 2 of the younger rats died within 24 h of injection. In addition, one animal (of 12 participants) was euthanized by protocol after the 5.6 mg/kg + Wheel Access condition (run before it was scheduled for 10 mg/kg + Wheel but after it completed 10 mg/kg No Wheel successfully); this individual was of the younger age group. Six of the 11 animals which were treated with 10 mg/kg in the No Wheel condition (all survived) were of the ∼40 wk old cohort. The one animal (of 7 treated) who died in the 10 mg/kg + Wheel Access at 23 °C ambient condition (Experiment 2) was 30 wks of age at the time. Thus it is unlikely that age or participation in prior drug studies had a major qualitative effect on the outcome. The most parsimonious interpretation is that wheel running poses additional risk in rats from ∼20 wk to ∼40 wk of age, regardless of prior drug history.

### 3.2 Wheel Activity

#### 3.2.1 Experiment 1

In the first experiment with ambient temperature under normal building control, but not explicitly monitored or altered, there were no significant effects of drug/wheel treatment condition, of time post-injection, nor of the interaction, on wheel rotations (Figure 1A). One animal’s wheel data after vehicle challenge was not collected due to equipment malfunction thus the analysis was between-groups for this measure.

**Figure 1.**
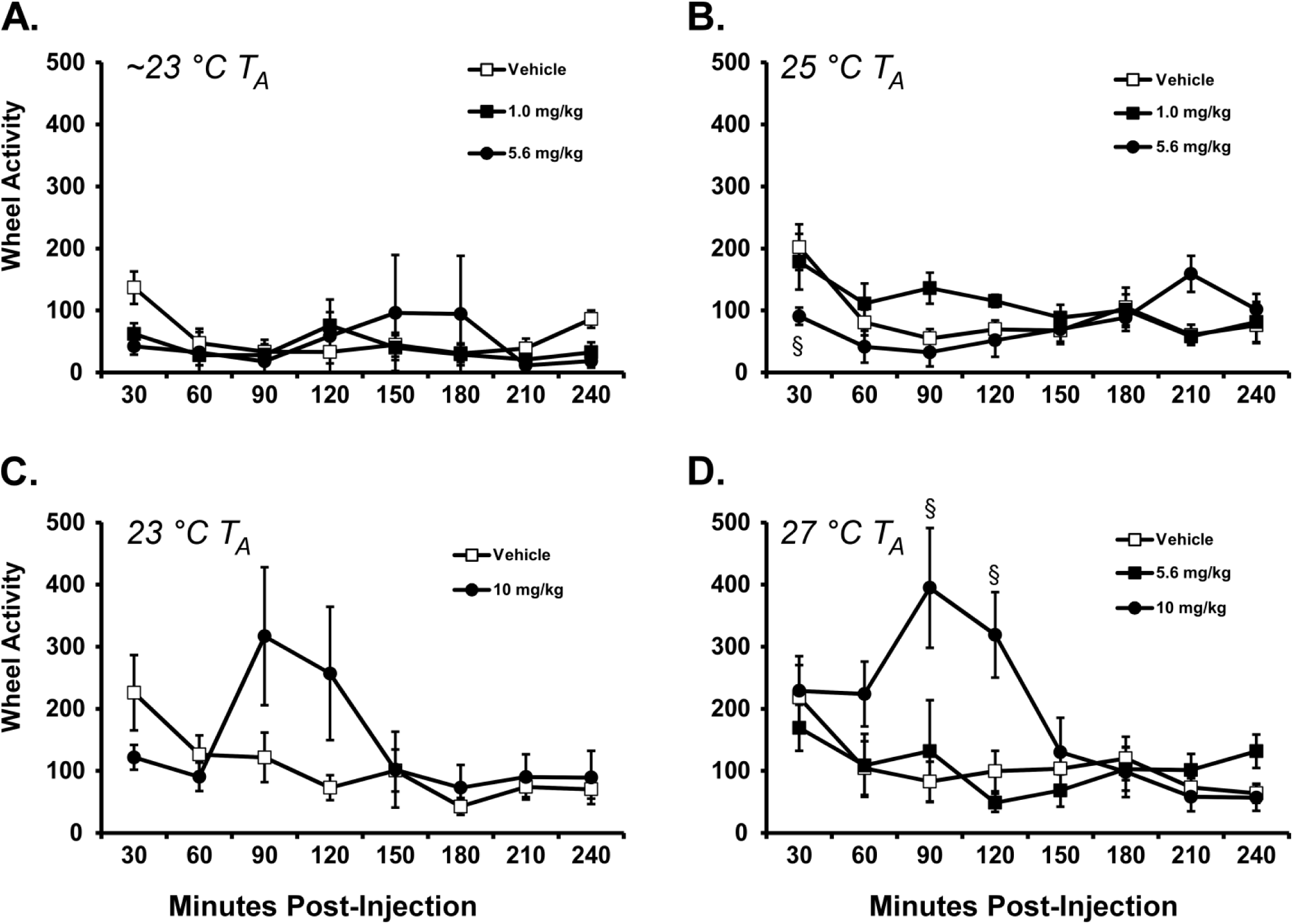
Mean wheel activity (average quarter rotations) for **A)**Rats (N = 5-6) following challenge with Vehicle or MDMA (5.6, 10 mg/kg, s.c.) under normal laboratory (∼23°C) T_A_ conditions; B) Rats (N = 7-8) following challenge with Vehicle or MDMA (5.6, 10 mg/kg, s.c.) under 25°C T_A_; C) challenge with Vehicle or 10 mg/kg MDMA, s.c., under 23°C T_A_; or D) Rats (N = 10-14) following challenge with Vehicle or MDMA (5.6, 10 mg/kg, s.c.) under 27°C T_A_. Significant differences between the highest MDMA dose and all other conditions at a given time post-injection is indicated by §. Bars indicate SEM.

#### 3.2.2 Experiment 2

A replication of the lower-dose experiment was then performed with the T_A_ controlled and monitored to 25°C in a group (N=8) of naïve male rats. The ANOVA confirmed that wheel activity was significantly changed (Figure 1B) by time post-injection [F (7, 119) = 6.03; P < 0.0001] and by the interaction of drug/wheel treatment condition with time post-injection [F (14, 119) = 4.12; P < 0.0001]. The post-hoc test further confirmed wheel activity was *lower* in the 5.6 mg/kg MDMA condition compared with the Vehicle (30 min post-injection) and 1.0 mg/kg MDMA (30, 90 min post-injection) conditions. The group was next challenged with vehicle or 10 mg/kg MDMA with T_A_ controlled at 23 °C (Figure 1C). One animal from the group was lost prior to this set of challenges for reasons unrelated to these experimental treatments, thus N=7 for most of the high-dose MDMA / 23°C T_A_ study. As noted under section 3.1, one of the remaining seven animals died after the 10 mg/kg MDMA Wheel access condition prior to participating in its last condition (10 mg/kg MDMA / No Wheel) thus analysis is between subjects. In this case the ANOVA confirmed that wheel activity was significantly changed across time post-injection [F (7, 84) = 2.98; P < 0.01] and determined by the interaction of drug/wheel treatment condition with time postinjection [F (7, 84) = 2.50; P < 0.05], however the post-hoc test did not confirm any significant differences in wheel activity between drug conditions at any *specific* times post-injection.

#### 3.2.3 Experiment 3

Out of the N=16 rats used in this study, N=14 completed the 5.6 mg/kg MDMA / No Wheel and the 10 mg/kg MDMA / Wheel conditions, N=13 completed the Vehicle / Wheel condition, N=11 completed the Vehicle / No Wheel, 5.6 mg/kg MDMA / Wheel and 10 mg/kg MDMA / No Wheel conditions due to the randomization of treatment conditions and subject mortality, as described in Section 3.1. Consequently the analysis treated drug/wheel condition as a between-groups factor since not all subjects were represented in all conditions. Wheel activity was again affected by drug treatment and time post-injection (Figure 1D) in this experiment and the ANOVA confirmed a significant effect of time post-injection [F (7, 175) = 6.06; P < 0.0001] and the interaction [F (7, 175) = 4.75; P < 0.0001] wheel activity. The post-hoc test confirmed that wheel activity was higher in the 10 mg/kg MDMA treatment compared with vehicle 90-120 min after injection.

Together the wheel data confirm and extend our prior results (Gilpin et al. 2011) by showing that wheel activity is suppressed or unchanged by doses of MDMA up to 5.6 mg/kg and increased by 10 mg/kg. The stimulant effect of 10 mg/kg MDMA on wheel activity was approximately equivalent from 23-27 °C T_A_.

### 3.3 Body Temperature

#### 3.3.1 Experiment 1

Body temperature was decreased by MDMA in a dose, time and wheel-access dependent manner (Figure 2A) as confirmed by a main effect of time post-injection [F (7, 35) = 4.76; P < 0.001) and of the interaction of drug/wheel treatment condition with time post-injection [F (35, 175) = 3.48; P < 0.0001) in the ANOVA. The post-hoc test confirmed temperature was *lower* in the 5.6 mg/kg MDMA / Wheel condition compared with all other conditions at 30 min post-injection and compared with all other conditions except the 5.6 mg/kg MDMA / No Wheel condition at 60 minutes post-injection.

**Figure 2.**
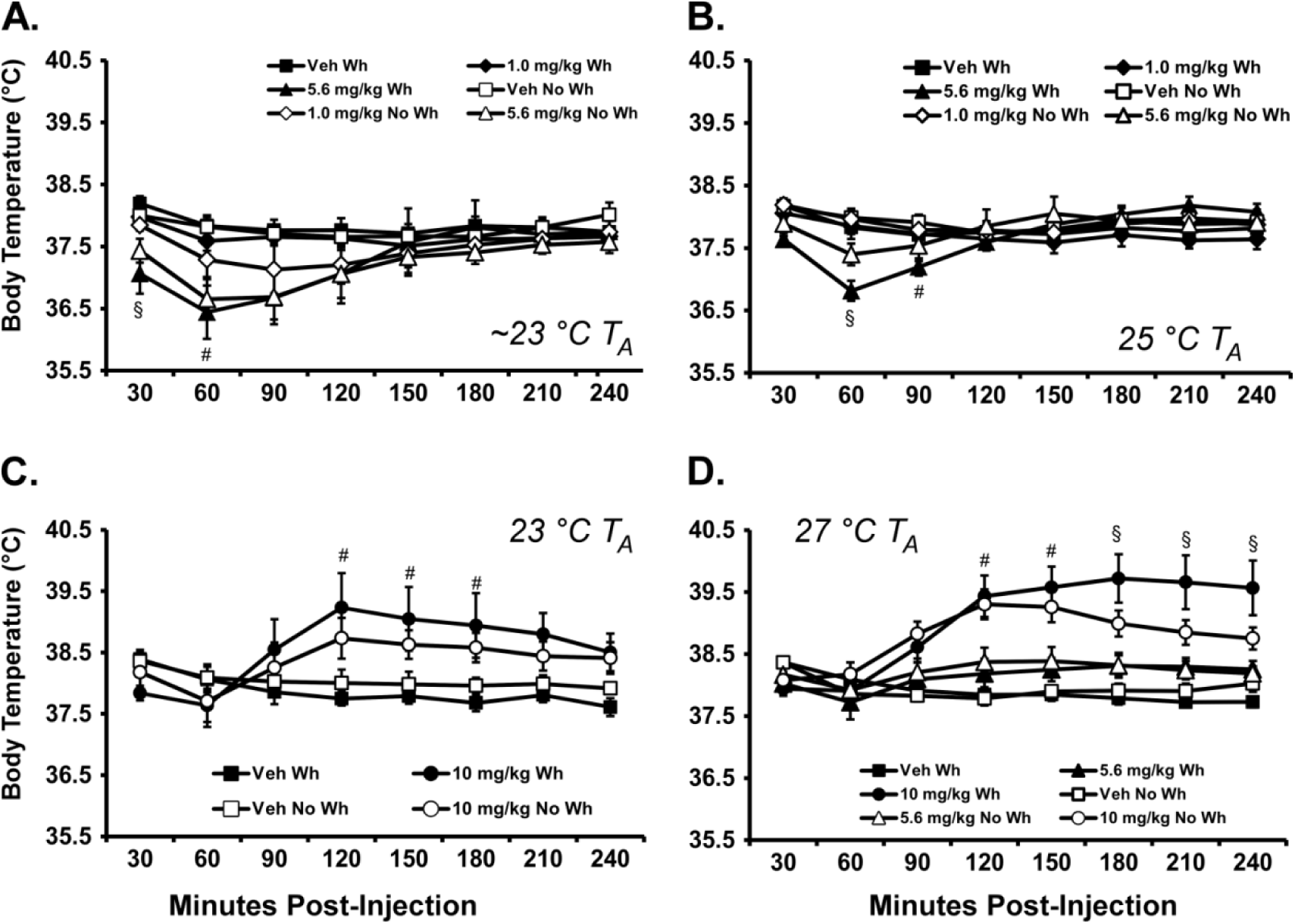
Mean temperature for **A)**Rats (N = 6) following challenge with 0, 5.6 or 10 mg/kg MDMA under normal laboratory (∼23°C) T_A_ conditions; B) Rats (N = 6-8) following challenge with 0, 5.6 or 10 mg/kg MDMA under 25°C T_A_; C) challenge with 0 or 10 mg/kg MDMA under 23°C T_A_; or D) Rats (N = 10-14) following challenge with 0, 5.6 or 10 mg/kg MDMA under 27°C T_A_. Drug treatments were repeated twice to incorporate conditions with and without wheel (**Wh**) access. Significant differences between the highest MDMA dose with Wheel access and all other conditions is indicated by §; and between the highest MDMA dose with Wheel and all other conditions except the corresponding highest MDMA / No-Wheel condition by #. Bars indicate SEM.

#### 3.3.2 Experiment 2

Body temperature was decreased by drug treatment under 25 °C T_A_ (Figure 2B) in this group and the ANOVA confirmed a main effect of time post-injection [F (7, 287) = 9.80; P < 0.0001] and of the interaction of drug/wheel treatment condition with time post-injection [F (35, 287) = 4.61; P < 0.0001]. The post-hoc test confirmed temperature was lower in the 5.6 mg/kg MDMA / Wheel condition compared with all other conditions at 60-90 min post-injection. (One animal’s temperature data was lost in the 1.0 mg/kgMDMA / Wheel condition due to technical error thus analysis was between-subjects.)

Body temperature was increased by 10 mg/kg MDMA under 23 °C T_A_ (Figure 2C) as confirmed by significant effects of time post-injection [F (7, 161) = 4.46; P < 0.0005] and of the interaction of drug/wheel treatment condition with time post-injection [F (21, 161) = 5.14; P < 0.0001] in the ANOVA.

The post-hoc test confirmed that temperature was higher after 10 mg/kg MDMA was administered with Wheel access compared with either Wheel (120-210 min post-injection) or No Wheel (120-180 min post-injection) vehicle conditions but not the 10 mg/kg MDMA / No Wheel condition.

#### 3.3.3 Experiment 3

The analysis treated drug/wheel condition as a between-groups factor since not all subjects were represented in all conditions, as outlined in Section 3.2.3, above. One individual’s 240 min post-injection value was a duplication of the 210 value in the 10mg/kg + Wheel condition due to a missing value (due to cooling the animal per protocol). The ANOVA confirmed a main effect of Drug/Wheel Condition [F (3, 45) = 10.67; P < 0.0001]; of time post-injection [F (7, 315) = 10.99; P < 0.0001] and the interaction [F (21, 315) = 12.23; P < 0.0001] on body temperature (Figure 2D). The post-hoc test confirmed that temperature was higher in the 10 mg/kg MDMA / Wheel condition compared with all other conditions from 180-240 min post-injection and compared with all other conditions except 10 mg/kg MDMA / No Wheel condition from 120-150 minutes post-injection.

### 3.4 Activity

#### 3.4.1 Experiment 1

There were no significant effects of drug treatment / wheel access condition or time post-injection on activity counts measured by the telemetry system in this experiment (Figure 3A).

**Figure 3.**
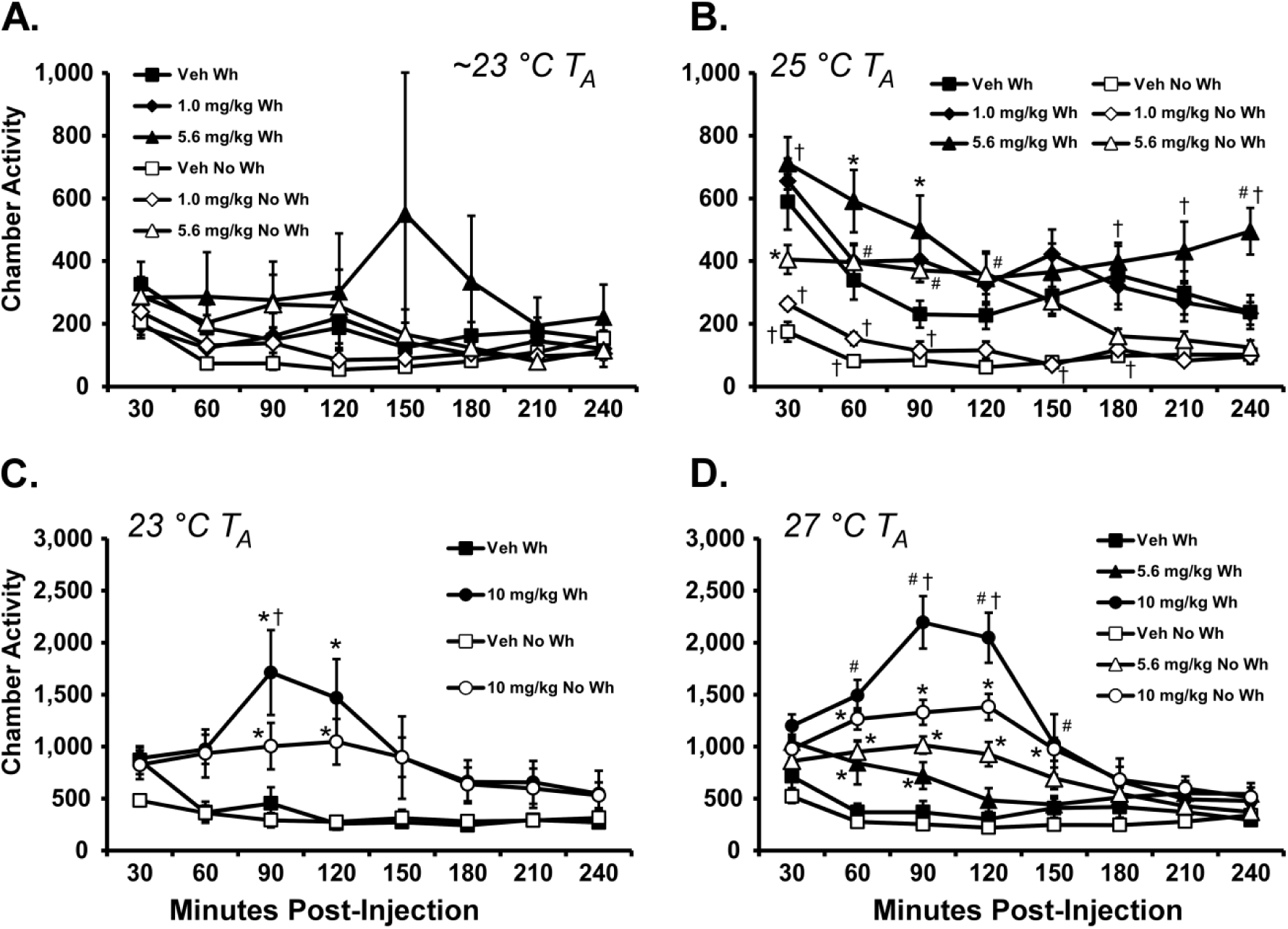
Mean chamber activity for **A)**Rats (N = 6) following challenge with 0, 5.6 or 10 mg/kg MDMA under normal laboratory (∼23°C) T_A_ conditions; B) Rats (N = 7-8) following challenge with 0, 5.6 or 10 mg/kg MDMA under 25°C T_A_; C) Rats (N = 6-7) challenge with 0 or 10 mg/kg MDMA under 23°C T_A_; or D) Rats (N = 10-14) following challenge with 0, 5.6 or 10 mg/kg MDMA under 27°C T_A_. Drug treatments were repeated twice to incorporate conditions with and without wheel (**Wh**) access. Note scale change between upper and lower panels. Significant differences from Vehicle (*) or Vehicle and 1.0 mg/kg (#) within Wheel access conditions are indicated. Significant differences between the Wheel access conditions within Veh/MDMA dose are indicated by f. Bars indicate SEM.

#### 3.4.2 Experiment 2

The activity in the home chamber was increased by both MDMA injection and wheel access (Figure 3B) when studies were conducted under 25°C T_A_. The ANOVA confirmed a main effect of Drug/Wheel Condition [F (5, 41) = 18.57; P < 0.0001], of time post-injection [F (7, 287) = 18.76; P < 0.0001] and the interaction [F (35, 287) = 2.15; P < 0.0005] on home chamber activity. The post-hoc test confirmed that activity was increased relative to the respective vehicle wheel condition after 5.6 mg/kg MDMA was administered either with (60-90, 240 minutes post-injection) or without (30-120 minutes postinjection) Wheel access. Likewise, activity differed between 1.0 and 5.6 mg/kg MDMA treatments in Wheel (240 minutes post-injection) and No Wheel (60-150 minutes post-injection) conditions. Home chamber activity was also higher in the Wheel vs No Wheel Conditions after Vehicle (30-60, 180 minutes post-injection), 1.0 mg/kg MDMA (30-90, 150 minutes post-injection) and 5.6 mg/kg MDMA (30, 180-240 minutes post-injection).

The ANOVA confirmed a Main effect of Drug/Wheel Condition [F (3, 23) = 9.67; P < 0.0005]; of time post-injection [F (7, 161) = 4.80; P < 0.0001] and the interaction [F (21, 161) = 2.09; P < 0.01] on home chamber activity (Figure 3C). The post-hoc test confirmed that activity was increased relative to the respective wheel condition vehicle 90-120 min after 10.0 mg/kg MDMA either with or without Wheel access. Activity was also higher 90 min after 10 mg/kg MDMA was administered in the Wheel access condition compared with the same dose without wheel access.

#### 3.4.3 Experiment 3

Chamber activity measured by telemetry was increased by both wheel access and MDMA administration under 27°C T_A_ (Figure 3D). The ANOVA confirmed a Main effect of Drug/Wheel Condition [F (5, 69) = 22.05; P < 0.0001]; of time post-injection [F (7, 483) = 24.05; P < 0.0001] and the interaction [F (35, 483) = 5.41; P < 0.0001] on home chamber activity. The Holm-Sidak post-hoc confirmed that activity was higher after 10 mg/kg MDMA compared with the respective vehicle treatment 60-150 min post-injection in the No Wheel and Wheel access conditions. The activity was also higher 90-120 min after injection of 10 mg/kg MDMA when the Wheel was available compared to the same dose in the No Wheel condition. In addition, the post-hoc test confirmed that activity was higher after 5.6 mg/kg MDMA compared with the respective vehicle condition when administered with Wheel access (60-90 min post-injection) or with No Wheel (60-120 min post-injection). Activity also differed significantly between 5.6 and 10 mg/kg MDMA treatments with Wheel access (60-150 min post-injection) but not under No Wheel conditions.

## 4. Discussion

This study examined the effect of wheel activity on the thermoregulatory effects of MDMA. The majority of prior studies of MDMA-induced activity in rodent models has reported *spontaneous* home chamber or open field activity changes after MDMA and, as such, could not make firm conclusions about the role of physical exertion. The manipulation of wheel access in the present study was successful in changing activity levels on the wheel and also in terms of chamber movement. The results show that higher levels of activity during the post-drug interval increases the magnitude of hyperthermia and the likelihood of fatality, particularly when a high dose is administered under ambient temperature (T_A_) that is thermoneutral for rats (Gordon 1990). Specifically, this study identified a threshold of approximately 27 °C to produce interactive effects of wheel activity to potentiate the hyperthermia caused by 10 mg/kg MDMA. Wheel access was also associated with an increase in 24-h mortality rate, consistent with our prior report (Gilpin et al. 2011). The 27 °C T_A_ threshold reported here for adverse effects of voluntary exercise is considerably lower than the 32 °C T_A_ reported to interact with *forced* running to exaggerate hyperthermia induced by 3 mg/kg MDMA (Tao et al. 2015). Additional work has shown that under 32 °C T_A_ 7.5 mg/kg MDMA, i.v., slightly increased the rate of gain, but not the eventual magnitude, of hyperthermia generated by forced treadmill activity in male rats (Zaretsky et al. 2015). The 32 °C T_A_ used in those two prior studies is likely to be above the thermoneutral range for rats (Gordon 1990) and is therefore less likely to generalize to real world conditions for humans (McNeill and Parsons) compared with the present investigation.

The fact that locomotor activity measured by the telemetry was higher in the wheel access versus the no wheel condition after 10 mg/kg MDMA (and 5.6 mg/kg MDMA under 25 °C T_A_) was unexpected since the *a priori* assumption was that time on the wheel would decrease time available for movement in the chamber. This result may potentially be attributed to an increase in locomotion due to repeated entry and exit from the wheel or perhaps a non-specific increase in general activity. For interpretive purposes within the present study, this outcome shows that physical activity of the rats during the wheel access condition may have increased even more than is reflected in the wheel rotations. Importantly, while wheel activity after 5.6 mg/kg MDMA at 25 °C T_A_ was less than in the related vehicle condition as in our prior work (Gilpin et al. 2011; Huang et al. 2012), chamber locomotor activity was greater. Overall, MDMA-associated increases in chamber activity were dose-dependent *within each wheel access condition*, further reinforcing the validity of the model.

In our prior study (Gilpin et al. 2011) 3/10 individuals died after 10 mg/kg MDMA at a ∼22°C ambient temperature. In two cases these individuals experienced 10 mg/kg with wheel access as the first condition, exhibited the highest temperature changes recorded along with high rates of running and died thereafter. In one case, the individual died after 10 mg/kg w/o wheel access. The present data further indicate that wheel access following 10 mg/kg MDMA enhances lethality, particularly as the ambient temperature increases. More precise delineation of lethality thresholds and mechanisms are not possible from this study, since it was not designed with death as an endpoint. A cooling protocol with humane euthanasia for animals that could not be stabilized was in place and thus experimental mortality was a mixture of euthanized animals, those that died before temperature could be normalized and individuals who were stable for hours but were nevertheless found dead overnight. These outcomes suggest that multiple mechanisms of fatality may result and justify further investigation into the role exercise plays in MDMA-associated fatality.

One minor limitation to the present results is the repeated measures design. Inevitably the fact that the experiments were conducted in groups of animals who differed in precise age and the number of prior drug challenges under various treatment conditions means some caution is warranted. Nevertheless, the results were internally consistent and were congruent with prior observations in many particulars. We previously showed that 5.6 mg/kg MDMA, i.p., suppresses wheel activity whereas 10.0 mg/kg increases wheel activity under normal laboratory T_A_ (∼22 °C) conditions (Gilpin et al. 2011), similar to Figure 1B. In that study only 10 mg/kg MDMA, i.p., caused a significant elevation of body temperature, consistent with the present effects. We have also shown that 5.6 mg/kg MDMA, s.c., increases telemetered home cage activity and body temperature under 30 °C T_A_ (Miller et al. 2013) which is here anticipated by the change from a hypothermia to a slight temperature increase cased by 5.6 mg/kg MDMA across the 25-27 °C T_A_ range (Figure 2B,D). Thus, the consistency with prior observations suggests that the repeated measures design of this investigation did not play a meaningful role in the outcome.

## 5. Conclusions

In conclusion this study demonstrates that spontaneous locomotor activity on a running wheel can increase the mean body temperature changes observed following the administration of MDMA. This was only the case for a dose that consistently elevated locomotor activity both in the home cage and on the wheel and when MDMA was administered at 27 °C ambient temperature. We did not observe this synergy of MDMA with wheel activity when administered at 23 °C or in our prior study of 10 mg/kg MDMA (Gilpin et al. 2011) administered at a similarly low ambient temperature. This study therefore provides the first experimental demonstration that physical exercise interacts with the effects of MDMA to produce adverse physiological and even lethal consequences.

## Contributors

M.A.T. designed the study which was implemented by research technical staff of the laboratory. Statistical analysis of the data, creation of figures and manuscript drafting was conducted by M.A.T.

## Acknowledgements

The author is grateful to Sophia A. Vandewater and Glen Dickinson for expert technical assistance. This is manuscript #29272 from The Scripps Research Institute.

